# Unsupervised genome-wide cluster analysis: nucleotide sequences of the omicron variant of SARS-CoV-2 are similar to sequences from early 2020

**DOI:** 10.1101/2021.12.29.474469

**Authors:** Georg Hahn, Sanghun Lee, Dmitry Prokopenko, Tanya Novak, Julian Hecker, Surender Khurana, Lindsey R. Baden, Adrienne G. Randolph, Scott T. Weiss, Christoph Lange

## Abstract

The GISAID database contains more than 1,000,000 SARS-CoV-2 genomes, including sequences of the recently discovered SARS-CoV-2 omicron variant and of prior SARS-CoV-2 strains that have been collected from patients around the world since the beginning of the pandemic. We applied unsupervised cluster analysis to the SARS-CoV-2 genomes, assessing their similarity at a genome-wide level based on the Jaccard index and principal component analysis. Our analysis results show that the omicron variant sequences are most similar to sequences that have been submitted early in the pandemic around January 2020. Furthermore, the omicron variants in GISAID are spread across the entire range of the first principal component, suggesting that the strain has been in circulation for some time. This observation supports a long-term infection hypothesis as the omicron strain origin.

## 1. Introduction

The emergence of new variants of the SARS-CoV-2 virus poses a great threat to the progress made by the ongoing vaccination efforts against COVID-19 worldwide (Dolgin, 2021). It is therefore important to identify and classify the newly emerging variants. In particular, we are interested in investigating how similar the nucleotide sequences of the newly emerging omicron variant (UCSC Genome Browser, 2021) are with respect to other nucleotide sequences that were collected from COVID-19 patients over the course of the pandemic.

In contrast to model-based approaches that assume an underlying phylogenetic tree-structure (Mousavizadeh and Ghasemi, 2021), a model-free, non-parametric way to assess the similarity of SARS-CoV-2 genomes at a genome-wide level is to utilize unsupervised cluster analysis based on the Jaccard similarity matrix and principal component analysis, as previously described by Hahn et al (2020a,b). This approach starts by aligning a given set of nucleotide sequences to a reference sequence, and subsequently translating all sequences into a Hamming matrix indicating all mismatches (mutations) with respect to the reference sequence. The Hamming matrix then serves as input to the Jaccard similarity measure/matrix, which results in a similarity index between 0 and 1 for all pairwise comparisons of sequences. Principal component analysis is then applied to the Jaccard similarity matrix to identify clusters and subgroups of SARS-CoV-2 genomes.

In this contribution, we apply the same methodology to calculate a similarity matrix between sequences of the SARS-CoV-2 virus observed since January 2020 worldwide, and newly added nucleotide sequences of the omicron variant. All results are based on sequences downloaded from the GISAID database (Elbe and Buckland-Merrett, 2017; Shu and McCauley, 2017). Using a principal component analysis, we observe that the sequences of the omicron variant are most similar to nucleotide sequences observed in January 2020, even though they are the newest/most recent sequences in our analysis. As the omicron variants do not form a cluster themselves, but are distributed over the entire range of the first principal component, our analysis provides support for the long-term infection hypothesis for the origin of the omicron strain (Chertow et al., 2021).

Due to its simplicity and computational speed, the applied unsupervised cluster analysis lends itself as a well-suited, assumption-free tool to continuously monitor data from public databases such as GISAID, with the aim to attempt to classify possibly emerging variants of interest for further follow-up analyses.

## 2. Methods

This section describes data acquisition and cleaning (Section 2.1), as well as details on the cluster analysis we perform (Section 2.2).

### 2.1 Data acquisition and cleaning

We base all findings reported in this communication on an image of all worldwide SARS-CoV-2 nucleotide sequences since 01 January 2020, available on the GISAID database (Elbe and Buckland-Merrett, 2017; Shu and McCauley, 2017) under the accession numbers in the range of EPI_ISL_403962 to EPI_ISL_13446977. These were downloaded on 24 June 2022.

We only take into account sequences satisfying all four data quality attributes offered by GISAID, that is *complete* (complete sequences are those of length at least 29,000bp), *high coverage* (defined as sequences with less than 1% N-bases), *with patient status* (defined as sequences for which meta information in the form of age, sex, and patient status is available), and *collection data complete* (defined as submissions with a complete year-month-day collection date). The metadata information we use is the geographic location in which each sequence was collected. This leaves 224,030 sequences for analysis. Since this number exceeds the size for which it is feasible to compute the Jaccard similarity matrix and subsequent principal component analysis, we further down-sample the dataset by taking an unbiased sample without replacement of size 10,000. The GISAID accession numbers of the 10,000 sequences we analyze are given in the supplementary material. Additionally, we save the accession numbers of all sequences of the omicron variant available on GISAID (those are 11,702 sequences), a proportion of which will be contained in our subsample of the entire database.

Next, we align all sequences to the official SARS-CoV-2 reference published on GISAID under the accession number EPI_ISL_402124. The alignment was performed with MAFFT (Katoh et al, 2002) using the *keeplength* option, and with all other parameters set to their default values. This allowed us to establish a well-defined window for comparison of length L=29891 base pairs.

### 2.2 Cluster analysis and regression analysis

Next, comparing each aligned nucleotide sequence to the reference genome allows us to compute a binary matrix X ∈ B^n x L^ in which an entry of X_ij_=1 indicates that the sequence with number i differs from the reference sequence at locus j, while an entry of 0 indicates no mismatch (here, the set B is defined as the binary numbers, that is B={0,1}). The number of rows n is taken as the number of nucleotide sequences we analyze. Therefore, X can be interpreted as a Hamming distance matrix on the genomes with respect to the reference genome.

As in Hahn et al (2020a,b) we proceed by computing the similarity of all pairs of sequences using the Jaccard similarity measure (Jaccard, 1901; Prokopenko et al, 2016; Schlauch et al, 2017) based on the matrix X. The Jaccard matrix J(X) has dimensions n times n, and each matrix entry (i,j) is a measure of similarity between the vector of mismatches for genome i and genome j. The computation of the Jaccard similarity measure was carried out with the R-package “locStra”, available on CRAN (Hahn et al, 2020c,d).

For visualization of the Jaccard similarity measures, we compute the first two principal components of the Jaccard matrix. We color the principal components based on (a) the geographic location that each sequence was collected in and (b) its time stamp (Elbe and Buckland-Merrett, 2017; Shu and McCauley, 2017), as well as (c) its clade (Hamed et al., 2021). Regarding the coloring by geographic location, we group all countries according to their WHO region, precisely AFRO (Regional Office for Africa), EMRO (Regional Office for the Eastern Mediterranean), EURO (Regional Office for Europe), SEARO (South-East Asia Region), WPRO (Western Pacific Region), and PAHO (Pan American Health Organization). Regarding the coloring by time stamp, we group all sequences by month (from January 2020 to June 2022) and look at their progression over time. For the coloring by clade, we differentiate according to the 11 clades of the SARS-CoV-2 genome available on GISAID (those are clades G, GH, GK, GR, GRA, GRY, GV, L, O, S, V).

We aim to quantify our claim that the aforementioned plot of the first two principal components of the Jaccard matrix indeed shows a progression of all nucleotide sequences in time. To this end, we perform a linear regression analysis in which the dependent variable (the Euclidean distance of each sequence from the origin in the principal component plot) is regressed on both the number of days since the start of the pandemic (defined as the number of days between 2020-01-01 and the time stamp of each sequence) and the sequence’s geographic origin (an ordinal variable with categories AFRO, EMRO, EURO, PAHO, SEARO, WPRO). For the geographic location, WPRO (Asia) is used as the reference factor level. The significance of the geographic location (the categorical variable) as a whole (as opposed to their respective categories returned by the “lm” command in R) is assessed via ANOVA analysis using the command “anova” in R.

## 3. Results

Figures 1-3 display the first two principal components of the Jaccard matrix, color coded by the WHO region each sequence was submitted from, by submission date, and by clade, respectively. In all three figures, the omicron sequences are depicted as triangles, while the other sequences are plotted as crosses.

**Figure 1.**
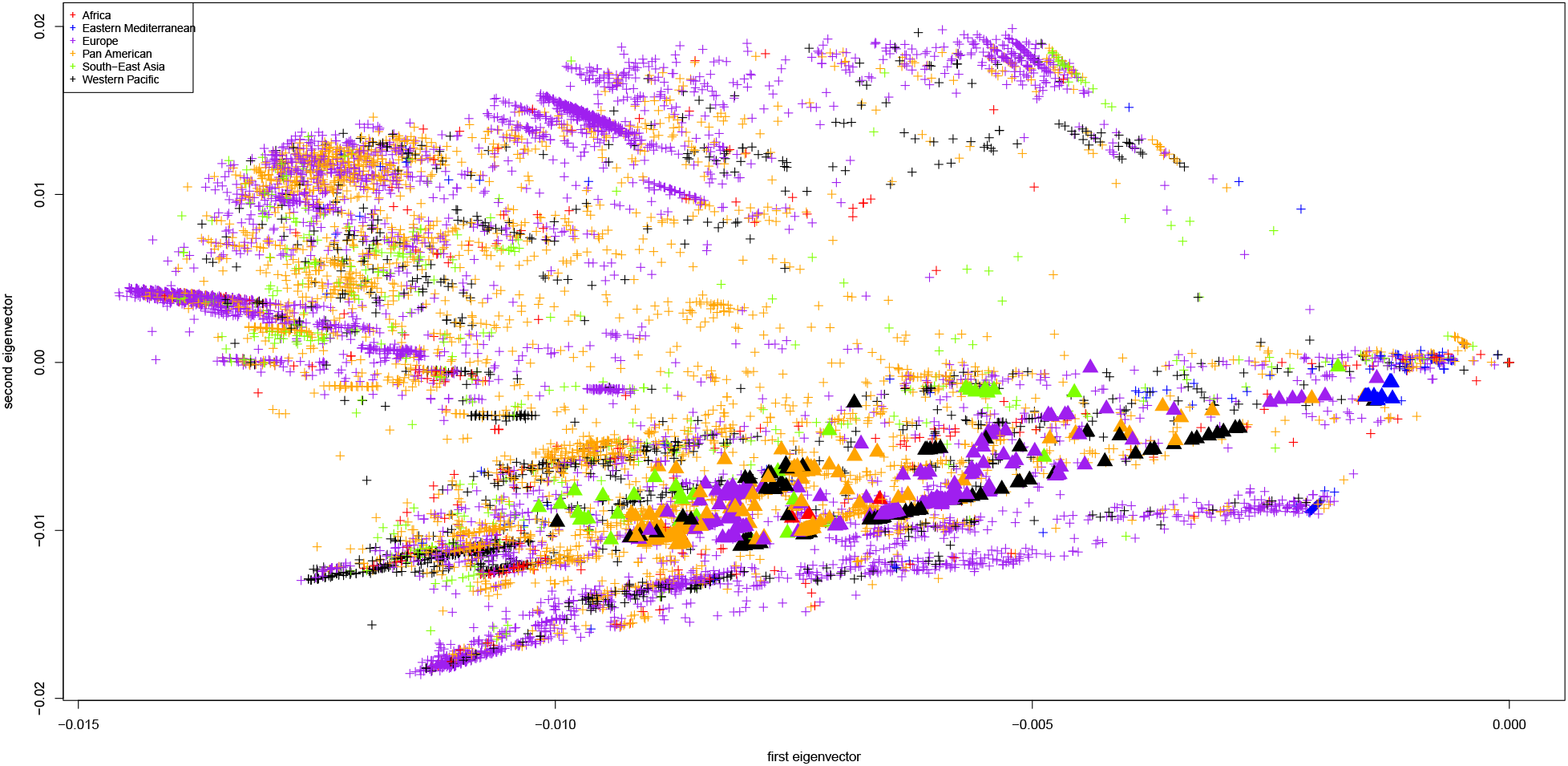
First two principal components of the Jaccard matrix, color coded by WHO region (AFRO in red, EMRO in blue, EURO in purple, PAHO in orange, SEARO in green, WPRO in black). Displayed are all sequences from GISAID, one point per sequence. The omicron samples are depicted as triangles.

**Figure 2.**
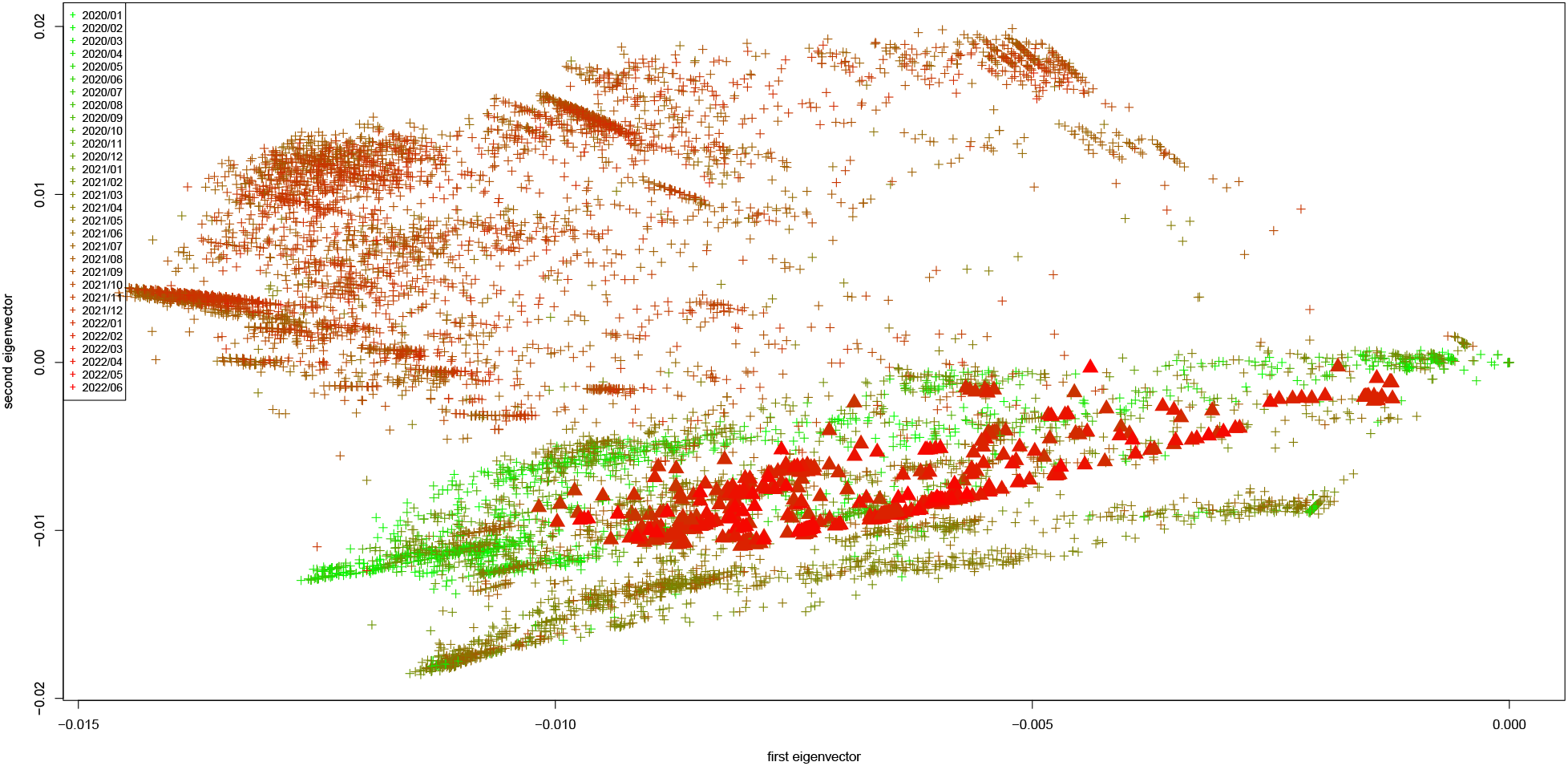
First two principal components of the Jaccard matrix, color coded by the time point each sequence was collected (from 01/2020 in green until 06/2022 in red). Displayed are all sequences from GISAID, one point per sequence. The omicron samples are depicted as triangles.

**Figure 3.**
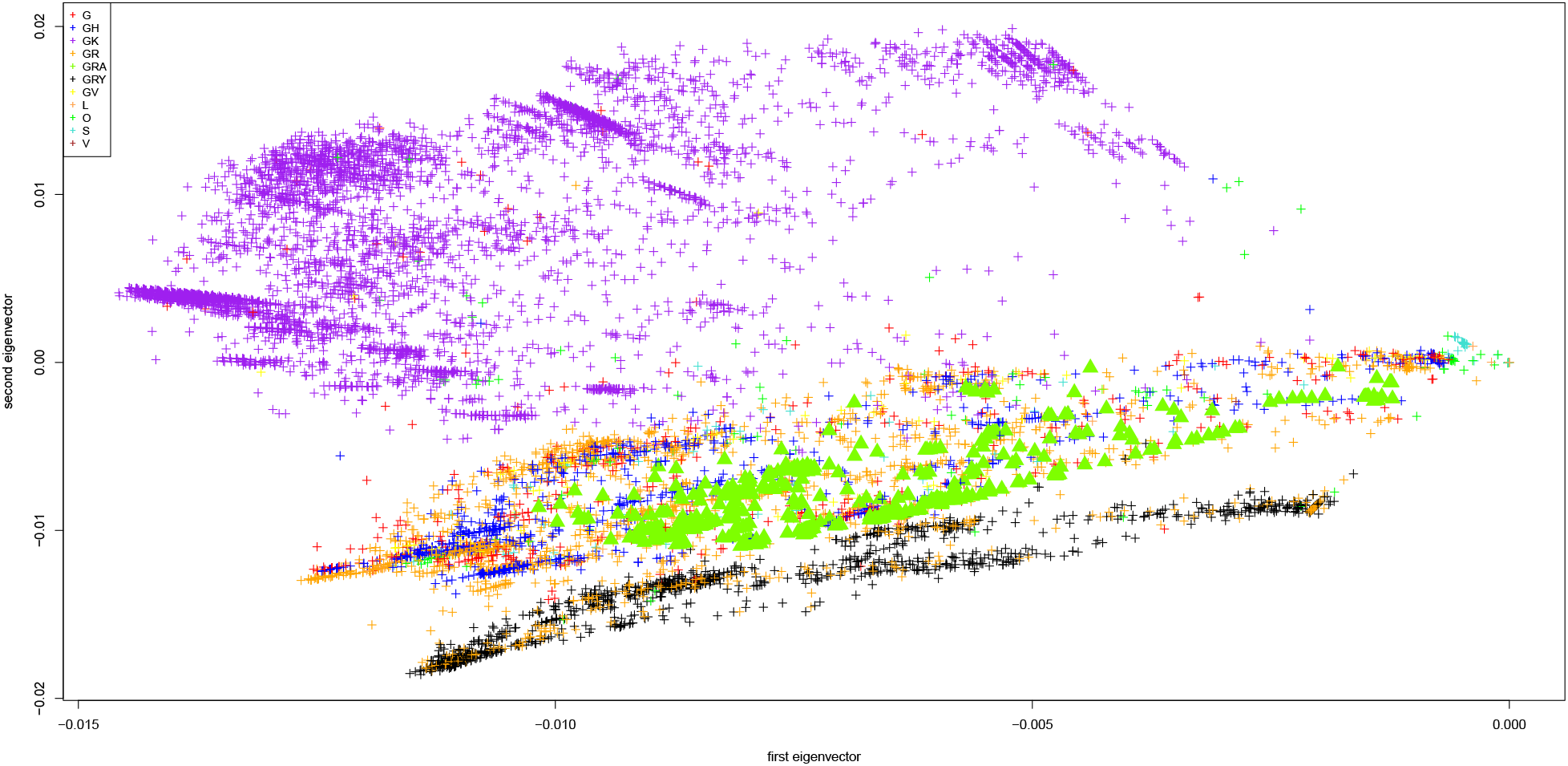
First two principal components of the Jaccard matrix, color coded by the clade of each sequence (clade G in red, GH in blue, GK in purple, GR in orange, GRA in green, GRY in black, GV in yellow, L in maroon, O in light green, S in turquoise, V in brown). Displayed are all sequences from GISAID, one point per sequence. The omicron samples are depicted as triangles.

As we have used the official SARS-CoV-2 reference genome for the alignment and the Hamming distance matrix assesses the magnitude of the deviation from the reference genome for all analyzed SARS-CoV-2 nucleotide sequences, the sequences that were submitted to GISAID from Wuhan in the early days of the pandemic are predominantly around the origin (0,0) of the plots. As in Hahn et al (2020b), we observe that the SARS-CoV-2 nucleotide sequences extend from the origin in a rather particular way, and form numerous clusters, distinctly according to their WHO region. For instance, we observe clusters of sequences from Europe in the upper left quadrant and upper parts of the plot, or from WPRO in the lower left quadrant. Interestingly, the sequences for the new omicron variant are somewhat outliers, with for instance the omicron cases observed in Europe (displayed in purple) being far off the European clusters, and closer to the origin. Moreover, the omicron sequences do not cluster themselves, which could hint at the fact that the strain has been in circulation for some time.

Similarly interesting as the clustering by geographic location is the coloring by time, displayed in Figure 2. We observe that the SARS-CoV-2 nucleotide sequences fall into three distinct point clouds, starting with the one in the center of the plot (for the sequences from 2020), after which sequences from early 2021 fall into the lower half of the plane. The sequences from late 2021 and from 2022 fall into the upper half of the principal component plot. Interestingly, the sequences stemming from the omicron variant, displayed as triangles, fall together with the older sequences from early 2020 in the similarity plot and are completely disconnected from the other sequences that have been submitted recently.

Finally, Figure 3 examines the clustering by clade. We again observe a distinctive clustering, with almost all of the GK clade (purple) clustering in the upper half of the plane, the GRY clade (black) falling almost exclusively into the lower half of the plane, and all other sequences being situated in the center of the plot. The omicron samples belong to the GRA clade (green) and, interestingly, do not cluster with the other GRA sequences.

Throughout the preparation of this manuscript, repeating the aforementioned analysis using earlier images of the GISAID database revealed that the clustering reported in Figures 1-3 is stable over time. This is noteworthy since all repeated analyses were carried out with independent, and thus entirely different, subsamples without replacement of size 10000.

The conclusion that the newly observed omicron samples are more similar to older samples stemming from early 2020 relies on the fact that Figure 2 indeed displays a progression in time. To support this claim analytically, we perform a linear regression as described in “Methods”. Table 1 provides the analysis results of the linear regression. The covariate for the submission date is highly significant (p-value < 2e-16), demonstrating that indeed the submission date is a significant predictor of the distance of each sequence from the origin.

**Table 1.**
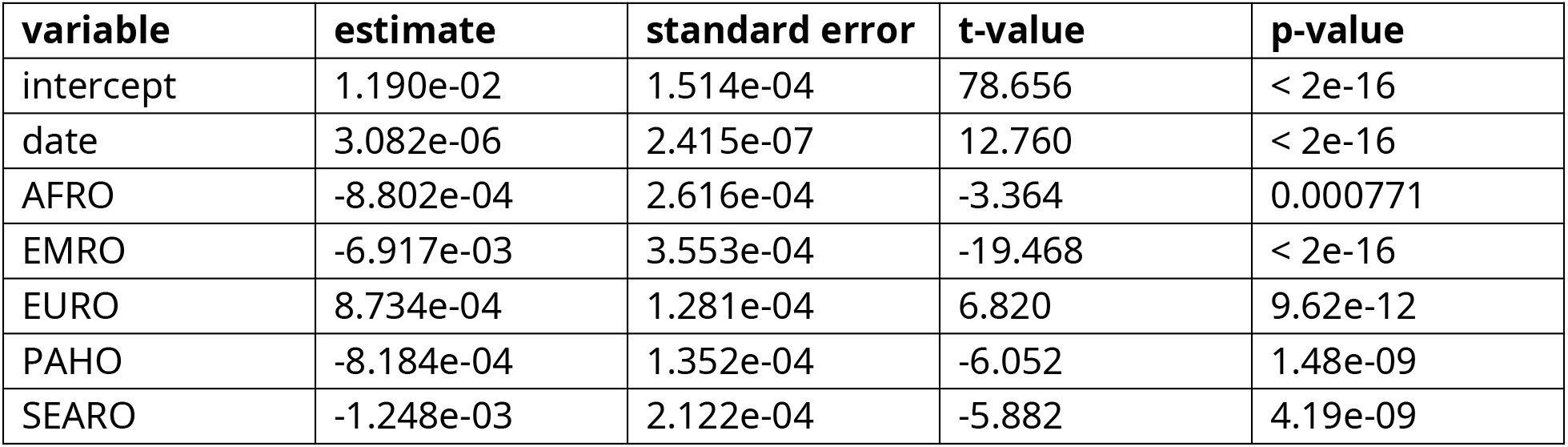
Results of a linear regression (with intercept) of the response variable “distance” (defined as the distance from the origin of each sequence in the principal component plot of Figure 2) on the date (defined as the number of days between the sequence’s time stamp and 2020-01-01) and the geographic location (categorical variable with categories AFRO, EMRO, EURO, PAHO, SEARO, WPRO, where WPRO for the Asian countries is used as the reference factor level). Using ANOVA analysis, the p-value for the categorical variable (the geographic location) as a whole is calculated as <2.2e-16.

| **variable** | **estimate** | **standard error** | **t-value** | **p-value** |
| --- | --- | --- | --- | --- |
| intercept | 1.190e-02 | 1.514e-04 | 78.656 | < 2e-16 |
| date | 3.082e-06 | 2.415e-07 | 12.760 | < 2e-16 |
| AFRO | -8.802e-04 | 2.616e-04 | -3.364 | 0.000771 |
| EMRO | -6.917e-03 | 3.553e-04 | -19.468 | < 2e-16 |
| EURO | 8.734e-04 | 1.281e-04 | 6.820 | 9.62e-12 |
| PAHO | -8.184e-04 | 1.352e-04 | -6.052 | 1.48e-09 |
| SEARO | -1.248e-03 | 2.122e-04 | -5.882 | 4.19e-09 |

## 4. Discussion

We performed a non-parametric principal component analysis on single stranded nucleotide sequences of the SARS-CoV-2 virus, which are publicly available in the GISAID database. We calculated a similarity score (the Jaccard measure), similarly to Hahn et al (2020b), with the aim to relate new cases of the so-called omicron variant of SARS-CoV-2 with the sequences that were collected during the past two years of the pandemic.

Using principal component plots, we observed that the new omicron sequences are closely related to sequences stemming from the early months of the pandemic, around January 2020, and are disconnected from the other sequences that have been submitted recently to GISAID, regardless of their origin. This could support the hypothesis that the omicron variant is linked to long-term infections. The wide spread of the omicron genomes with respect to the first principal component could suggest that the omicron strain has been in circulation for some time.

## References

Dolgin, E. (2021). Omicron is supercharging the COVID vaccine booster debate. Nature, https://doi.org/10.1038/d41586-021-03592-2

Chertow D., Stein S., Ramelli S., Grazioli A., Chung J.-Y., Singh M., Yinda C.K., Winkler C., Dickey J., Ylaya K., Ko S.H., Platt A., Burbelo P. Quezado M., Pittaluga S., Purcell M., Munster V., Belinky F., Ramos-Benitez M., Boritz E., Herr D., Rabin J., Saharia K., Madathil R., Tabatabai A., Soherwardi S., McCurdy M., Peterson K., Cohen J., de Wit E., Vannella K., Hewitt S., and Kleiner D. (2021). SARS-CoV-2 infection and persistence throughout the human body and brain. https://doi.org/10.21203/rs.3.rs-1139035/v1

Elbe S. and Buckland-Merrett G. (2017). Data, disease and diplomacy: GISAID’s innovative contribution to global health. Global Challenges, 1:33–46.

Hahn G., Lee S., Weiss S.T., and Lange C. (2020). Unsupervised cluster analysis of SARS-CoV-2 genomes indicates that recent (June 2020) cases in Beijing are from a genetic subgroup that consists of mostly European and South(east) Asian samples, of which the latter are the most recent. bioRxiv, pages 1–8, https://doi.org/10.1101/2020.06.22.165936

Hahn G., Lee S., Weiss S.T., and Lange C. (2020). Unsupervised cluster analysis of SARS-CoV-2 genomes reflects its geographic progression and identifies distinct genetic subgroups of SARS-CoV-2 virus. Genet Epidemiol, 45(3):316–323.

Hahn G., Lutz S.M., Hecker J., Prokopenko D., Cho M.H., Silverman E., Weiss S.T., and Lange C. (2020). locstra: Fast analysis of regional/global stratification in whole genome sequencing (wgs) studies. Genet Epidemiol, 45(1):82–98.

Hahn G., Lutz S.M., and Lange C. (2020). locStra: Fast Implementation of (Local) Population Stratification Methods (v1.3). https://cran.r-project.org/package=locStra.

Hamed S.M., Elkhatib W.F., Khairalla A.S., and Noreddin A.M. (2021). Global dynamics of SARS-CoV-2 clades and their relation to COVID-19 epidemiology. Sci Rep 11, 8435 (2021).

Jaccard P. (1901). Étude comparative de la distribution florale dans une portion des Alpes et des Jura. Bull Soc Vaud Des Sci Nat, 37:547–579.

Katoh K., Misawa K., Kuma K., and Miyata T. (2002). MAFFT: a novel method for rapid multiple sequence alignment based on fast Fourier transform. Nucleic Acids Research, 30(14): 3059–3066.

Mousavizadeh L. and Ghasemi S. (2021). Genotype and phenotype of COVID-19: Their roles in pathogenesis. Journal of Microbiology, Immunology and Infection, 54(2):159–163.

Prokopenko D., Hecker J., Silverman E., Pagano M., Nöthen M., Dina C., Lange C., and Fier H. (2016). Utilizing the Jaccard index to reveal population stratification in sequencing data: a simulation study and an application to the 1000 Genomes Project. Bioinformatics, 32(9):1366–1372.

Schlauch D., Fier H., and Lange C. (2017). Identification of genetic outliers due to sub-structure and cryptic relationships. Bioinformatics, 33(13):1972–1979.

Shu Y. and McCauley J. (2017). GISAID: Global initiative on sharing all influenza data -- from vision to reality. EuroSurveillance, 22(13):30494.

UCSC Genome Browser on SARS-CoV-2 (2021). Omicron variant: https://genome.ucsc.edu/cgi-bin/hgTracks?hgsid=1237196085_IsfCVVz6HLtTQ0q0pmGkwwWhAaWH&db=wuhCor1&position=lastDbPos

